# Large-scale algorithmic search identifies stiff and sloppy dimensions in synaptic architectures consistent with murine neocortical wiring

**DOI:** 10.1101/2021.11.13.468127

**Authors:** Tarek Jabri, Jason N. MacLean

**Affiliations:** Department of Neurobiology, The University of Chicago, Chicago, Illinois, United States of America; Committee on Computational Neuroscience, The University of Chicago, Chicago, Illinois, United States of America; Neuroscience Institute, Chicago, Illinois, United States of America

## Abstract

Complex systems can be defined by “sloppy” dimensions, meaning that their behavior is unmodified by large changes to specific parameter combinations, and “stiff” dimensions whose change results in considerable behavioral modification. In the neocortex, sloppiness in synaptic architectures would be crucial to allow for the maintenance of asynchronous irregular spiking dynamics with low firing rates despite a diversity of inputs, states, and both short- and long-term plasticity. Using simulations on neural networks with first-order spiking statistics matched to firing in murine visual cortex while varying connectivity parameters, we determined the stiff and sloppy parameters of synaptic architectures across three classes of input (brief, continuous, and cyclical). Algorithmically-generated connectivity parameter values drawn from a large portion of the parameter space reveal that specific combinations of excitatory and inhibitory connectivity are stiff and that all other architectural details are sloppy. Stiff dimensions are consistent across input classes with self-sustaining synaptic architectures following brief input occupying a smaller subspace as compared to the other input classes. Experimentally estimated connectivity probabilities from mouse visual cortex are consistent with the connectivity correlations found and fall in the same region of the parameter space as architectures identified algorithmically. This suggests that simple statistical descriptions of spiking dynamics are a sufficient and parsimonious description of neocortical activity when examining structure-function relationships at the mesoscopic scale. Additionally, coarse graining cell types does not prevent the generation of accurate, informative, and interpretable models underlying simple spiking activity. This unbiased investigation provides further evidence of the importance of the interrelationship of excitatory and inhibitory connectivity to establish and maintain stable spiking dynamical regimes in the neocortex.

**Author Summary:** Connections between neurons are continuously changing to allow learning and adaptation to new stimuli. However, the ability of neural networks to vary these connections while avoiding excessively high- or low-activity states is still not well understood. We tackled this question by studying how changes in the parameters of connectivity within and between different neuronal populations impacted network activity in computational models. We identified specific combinations of parameters, deemed “stiff”, that must be maintained to observe activity consistent with recordings from murine visual cortex, while the rest of the parameters can be varied freely with minimal effects on activity. Our results agree with experimentally measured connectivity statistics demonstrating the importance of balancing opposing forces to maintain activity in a natural regime.

## Introduction

Local synaptic connectivity in neocortex is fundamental to the generation and stipulation of the spiking dynamics [1,2] that underlie the formation of percepts, decisions, and the generation of appropriate behavioral responses. However, the rules that govern the mesoscopic-scale relationships between local synaptic architectures and spiking activity remain unclear. On one hand, synaptic architectures must be highly dynamic since they underlie, at least in part, the storage of information [3] and generate the range of spiking activity corresponding to distinct brain states [4]. On the other hand, it is clear that aberrant synaptic wiring can give rise to detrimental spiking behaviors and pathophysiology such as epilepsy [5].

Modeling and theoretical analysis are essential complements to experimental investigations of structure-function relationships since they enable precise manipulation of simulated connectivity [6,7]. Any model of a biological system involves a large number of free parameters that cannot always be determined from experimental data or that vary greatly from one observation to the other. Nevertheless, numerous models of neural systems have successfully replicated key aspects of neuronal network dynamics including different activity patterns [8–10] and receptive field properties [11–13]. Given the difficulty of estimating the exact values of connectivity parameters and given the variability in these values observed *in-vivo*, a reasonable hypothesis for why these models work is that multiple combinations of parameters can result in similar network activity [14]. Together, these observations are indicative of sloppy systems, whose behavior depends only on a few stiff combinations of parameters while the majority of parameters are not critical for accurate predictions of the system’s behavior [15].

Sloppiness is a universal feature of models in systems biology [6,15,16]. For example, ionic conductances within individual neurons have consistently been found to vary greatly across neurons and between individuals despite regularity in spiking activity [17–19]. At the neuronal circuit level, stability and state changes are mediated by a subset of neurons described by a small number of stiff parameter combinations while the parameters of the remainder of the neurons are sloppy [20,21].

What remains unclear, however, are the stiff and sloppy parameter combinations that define synaptic architectures capable of producing spiking that is low rate, irregular, and asynchronous—the dominant dynamical regime of the neocortex [1,22–24]. The conservation of both connectivity and wiring cost across different species [25] is a strong incentive to find the stiff and sloppy dimensions of synaptic architectures. Moreover, delineating these dimensions of neocortical wiring can better constrain cortical models. Understanding the stiff and sloppy dimensions also carries implications beyond the realm of organic neural systems into artificial neural networks (ANNs). It has been shown that connectivity patterns are directly related to the dimensionality of the activity in recurrent spiking neural networks (SNNs), with the latter decreasing as overall connectivity increases [26]. Hence, delineating the small number of parameters that describe stiff dimensions of network connectivity will allow further studies to capture meaningful functional and computational principles that define those networks.

Here, we survey a large portion of the synaptic connectivity parameter space to create wiring diagrams and identify parameter combinations capable of producing activity matched to murine visual cortex (V1). We identify the stiff and sloppy parameter combinations of synaptic architectures responsible for producing naturalistic activity and compare these algorithmically identified connectivity parameters to recent experimental values finding them to be largely in agreement.

## Results

### Grid search for synaptic architectures producing naturalistic spiking

In order to evaluate the impact of specific synaptic connectivity parameter combinations on the statistics of spiking, we carried out large scale simulations of spiking neural network (SNN) models. SNNs were composed of both excitatory and inhibitory adapting exponential leaky integrate-and-fire neurons (AdEx) [27] connected with conductance-based synapses (Fig 1A; see methods). In previous work, we have shown that conductance-based synapses are crucial to accurately simulate neuronal integration of synaptic inputs—a critical consideration when evaluating structure-function hypotheses [8,28]. We conducted a grid search over a range of synaptic connectivity parameters defined by both excitatory and inhibitory connectivity and then quantified the outcome in SNN model behavior as connectivity parameter values changed (Fig 1B). Specifically, we determined which connectivity parameter combinations were capable of producing sustained spiking activity matched to *in vivo* spiking of murine visual cortex [29–33]. The performance of the networks was quantified using a set of objective functions, each of which corresponded to individual first-order statistical descriptors of spiking activity: firing rates, synchrony, and fraction of trials with sustained activity (Fig 1C; see methods).

**Fig 1.**
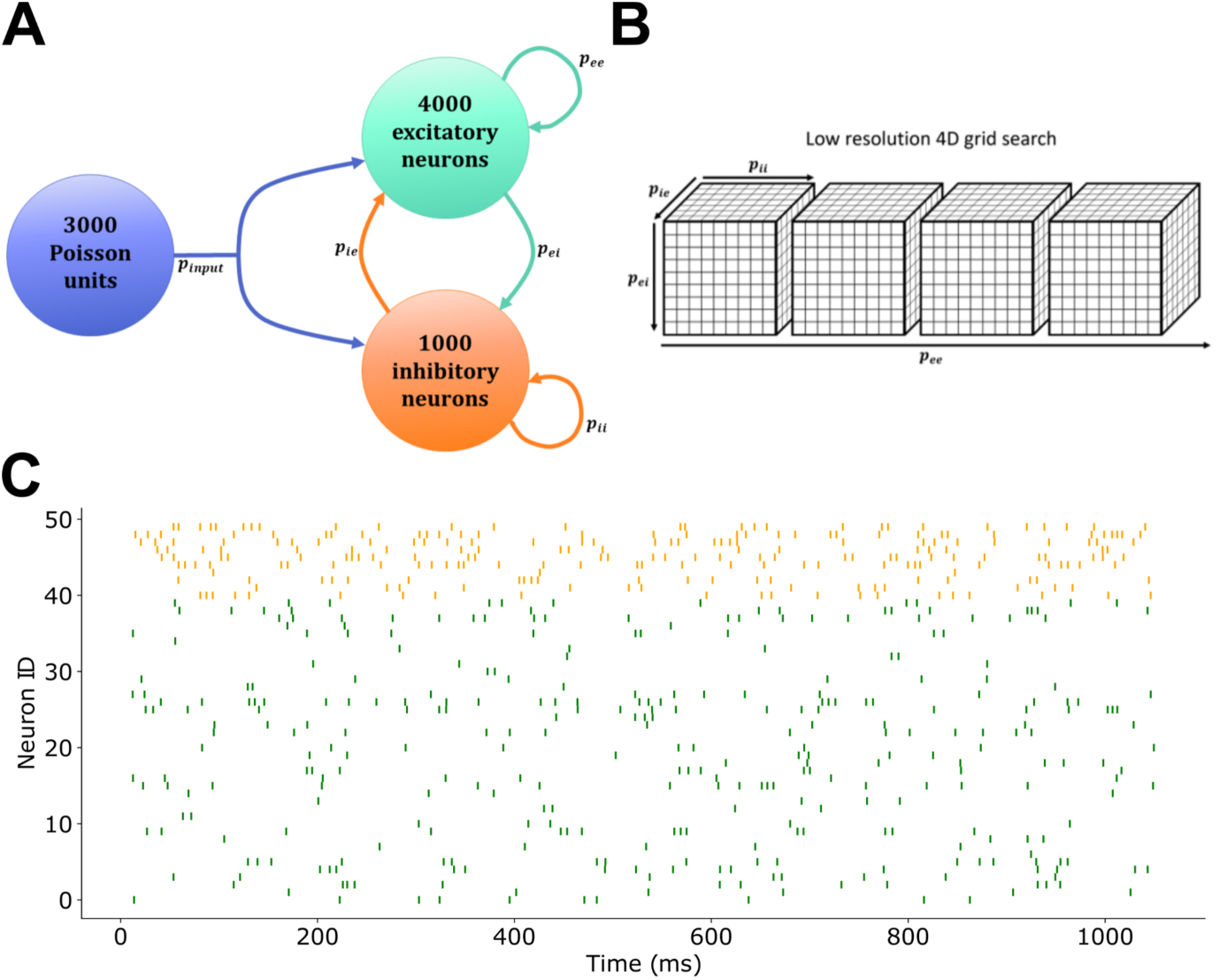
Network structure and activity. (A) Simulated network composition. (B) Design of grid search of synaptic architectures. (C) Raster plot showing the timing of spikes of 50 example neurons, for ease of presentation, in the network with connectivity probabilities that gave the lowest firing rates in the target range using continuous input. Neurons 0-39 (green) are excitatory and neurons 40-49 (orange) are inhibitory.

We began by identifying combinations of parameters which produced firing rates between 8 and 15 Hz [33] and synchrony scores corresponding to a Van Rossum distance greater than 4 in response to the three classes of input (constant **continuous, cyclical** continuous, **brief**). Notably, the input classes fall along a continuum of durations and as the inputs become increasingly brief, increased emphasis is placed on network architectures capable of producing self-sustaining activity following input. The networks were first tested using continuous input from Poisson units firing with rates drawn from a log-normal distribution with a mean of 17±5.3 Hz. Out of the 14461 unique connectivity parameter combinations tested, 5080 resulted in sustained activity for the length of the simulation more than 50% of the time. Of those, 579 synaptic wiring diagrams showed average excitatory firing rates in the desired range 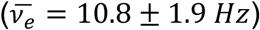. Those networks had a mean synchrony score of 1.03±0.06 using the fast synchrony measure (see methods; 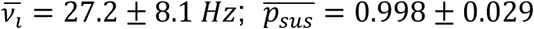). Networks receiving cyclical input (1.67Hz with maximal firing rates of units drawn from the same log-normal distribution) showed a lower number of successful parameter combinations at 199, including 47 networks with rates also between 8 and 15 Hz 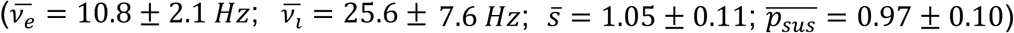. Finally, we evaluated architectures capable of producing self-sustaining activity in response to brief (300ms) excitatory Poisson input. We similarly simulated 73205 trials for 14461 synaptic architectures corresponding to a range of different parameter combinations. Self-sustained activity is the hardest to achieve, resulting in the lowest number of successful networks: 241 trials resulted in self-sustained activity, which in turn corresponded to 44 unique parameter combinations that produced self-sustained activity in at least 50% of the simulations. Of these 44 networks, 25 also scored well on the other objective functions and exhibited an average firing rate of excitatory neurons between 8 and 15 Hz (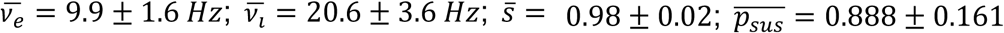; Fig 2B). Grid search resolution was low, in order to evaluate large ranges of parameter combinations for synaptic architectures. Thus, the low number of viable synaptic wiring diagrams should not be interpreted as indicative of a scarcity of viable architectures. In fact, subsequent calculations of the FIM (discussed below) show that within a narrow range (±0.01) around the parameter combinations found, all networks exhibit sustained activity with low firing rates and low synchrony levels.

**Fig 2.**
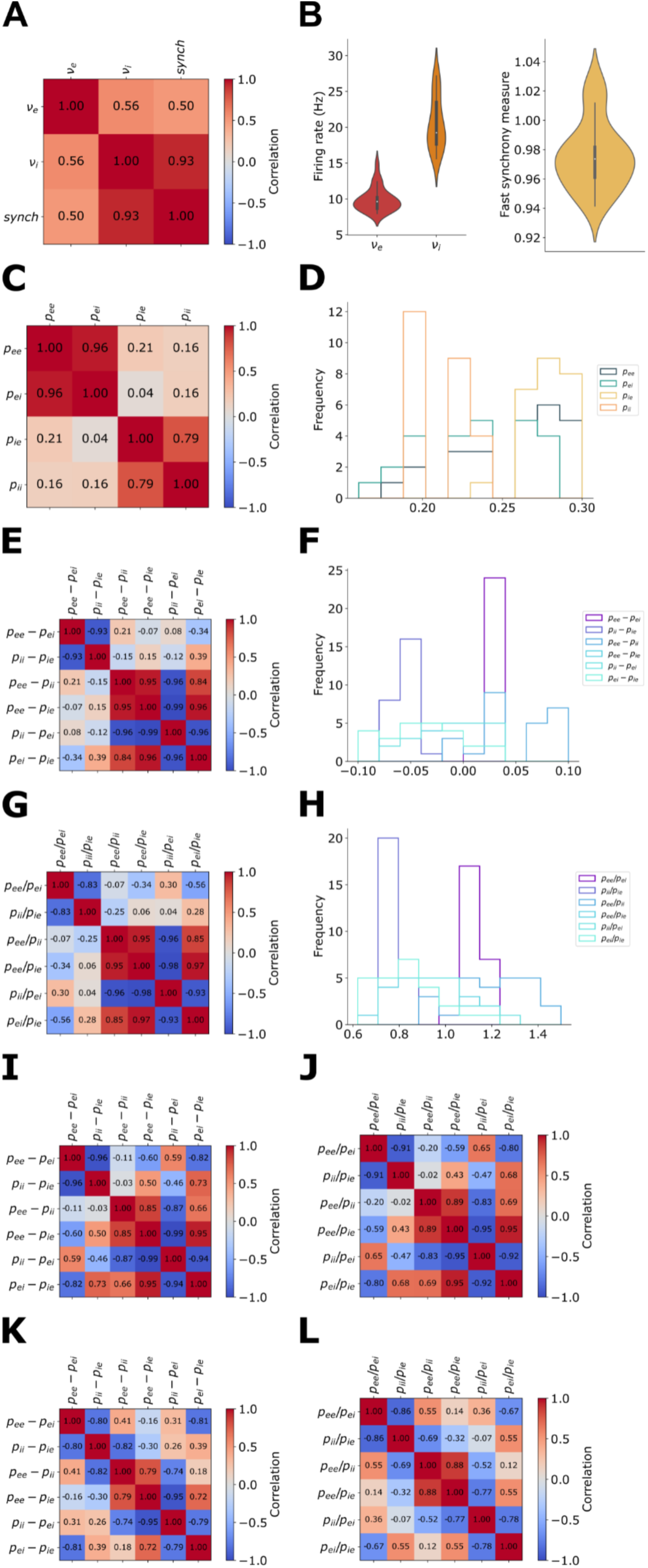
Analysis of networks from grid search with rate-matched sustained activity shows correlations between classes of connection. (A) Correlation values between spiking measures of networks using brief input. (B) Distribution of excitatory and inhibitory firing rates (*v*_*e*_ and *v*_*i*_) and synchrony measures of networks following brief input. (C) Pairwise Pearson correlation coefficients between the four connectivity parameters of those successful networks following brief input. (D) Distribution of the four parameters of connectivity in successful networks using brief input. (E)&(I)&(K) Pairwise Pearson correlation coefficients between differences of connectivity probabilities using brief, cyclical, and continuous inputs respectively. (F) Distribution of differences between connectivity probabilities from networks using brief input. (G)&(J)&(L) Pairwise Pearson correlation coefficients between ratios of connectivity probabilities using brief, cyclical, and continuous inputs respectively. (H) Distribution of differences between connectivity probabilities from networks using brief input. All networks considered here sustained their activity in more than 50% of the trials and had excitatory firing rates between 8 and 15 Hz.

This limited number of possible networks spread out over the entire range of parameters tested (Fig 2D) is, by definition, indicative of both sloppiness and stiffness of spiking neural networks. For the rest of the analyses, we will only consider networks that achieved sustained activity for each type of input.

It is noteworthy that despite the fact that the synchrony level was calculated using only excitatory spikes (see methods), the synchrony score is highly correlated with the inhibitory firing rate but not the excitatory one (Fig 2A). This confirms that this measure of synchrony does not simply scale with the number of spikes. Moreover, it is consistent with the role of local inhibition setting spike timing in the excitatory pool [34].

### Correlated components of naturally spiking synaptic architectures

As an initial investigation of potentially stiff parameter combinations, we examined the differences and ratios between pairs of connectivity probabilities for the parameter combinations that resulted in sustained activity. We observed that certain differences in the connection likelihoods were highly correlated with each other (either positively or negatively), while others were uncorrelated (Figs 2E and 2G). Specifically, *p*_*ee*_ – *p*_*ie*_ and *p*_*ei*_ − *p*_*ie*_ were strongly and positively correlated at 0.96 in the case of brief input, while *p*_*ee*_ − *p*_*ie*_ and *p*_*ii*_ − *p*_*ei*_ consistently showed the strongest negative correlations (−0.99 for brief and cyclical inputs, -0.95 for continuous inputs; Figs 2E, 2I, and 2K). The same pattern of correlations was observed among pairs of ratios (e.g. *p*_*ee*_/*p*_*ie*_ and *p*_*ee*_/*p*_*ii*_ very strongly correlated at 0.95, 0.89, 0.88 in the case of brief, cyclical, and continuous input respectively; Figs 2G, 2J, and 2L) indicating that networks that result in sustained activity are more likely to have connectivity parameters that follow these linear relationships between certain differences and ratios of excitatory and inhibitory connectivity. In other words, certain combinations of differences or combinations of ratios may constitute stiff parameters, while the others may be sloppy. In fact, especially in the case of networks receiving brief input, when looking at the pairs of differences that are highly correlated, it appeared that the majority of the networks had the same values for these differences despite having very different probabilities of connectivity (Figs 2F and 2H). Similar strong correlations were found among differences and ratios of wiring parameters in networks sustaining their activity with low firing rates using both brief and cyclical inputs (Figs 2I and 2J). When not restricting the firing rate to the target range the correlations remained but with lower magnitudes. Networks that exhibited sustained low-rate activity in response to continuous input (n=579) also exhibited the majority of the same significant correlations (Figs 2K and 2L). We note that if we did not control for firing rate, these correlations were less apparent in networks that spiked in response to continuous inputs. To confirm that these correlations that appear for rate-matched networks are not statistical artifacts due to the relatively small number of networks considered (∼11% of networks with sustained activity in more than 50% of the trials), we matched the sample size (n=579) in different sets of networks randomly selected from those that showed sustained activity in more than 50% of the trials and evaluated the correlations among the pairs of parameters. The lack of strong correlations in the case of randomly selected networks confirms that the results that we observed are related to the rates of those networks being in the target range.

These correlated parameter combinations are additional indicators of the presence of stiff dimensions within synaptic architectural parameter combinations when matching stable spiking activity in the network to *in vivo* recordings. It is notable that despite the fact that different networks showed sustained spiking with each type of input, the same pairs of connectivity parameter combinations were crucial to the production of sustained and murine-matched activity regardless of input type.

### Experimentally measured synaptic wiring in mouse visual cortex agrees with algorithmically-identified correlations

To evaluate the biological plausibility of the networks that we identified using grid search, we compared the algorithmically generated wiring of the SNNs to the values of measured connectivity probabilities recently reported by the Allen Institute for Brain Science [32]. To match the parameters varied in the grid searches, it was necessary to coarse grain the reported connection likelihoods from all classes of inhibitory interneurons and excitatory neurons regardless of laminar location by summarizing all of the connectivity measures as four probability values for excitatory and inhibitory neurons (Fig 3A; see methods). We also coarse grained the reported connectivities according to laminar position allowing us to separately compare individual layers 2/3, 4, and 5 with the algorithmically identified architectures. For these simulations, input connectivity probability was maintained at 10% for the entire cortex and for L4, but otherwise was based on the reported connectivity probabilities from L4 for layers 2/3 and layers 2/3 to L5 [32]. Despite the fact that after coarse graining some of the resulting connectivity probabilities were not in the range tested in the grid search and despite significant differences in connectivity probabilities between the different lamina, all of the networks exhibited sustained activity in response to continuous input consistent with the initial study (Fig 3B) [32].

**Fig 3.**
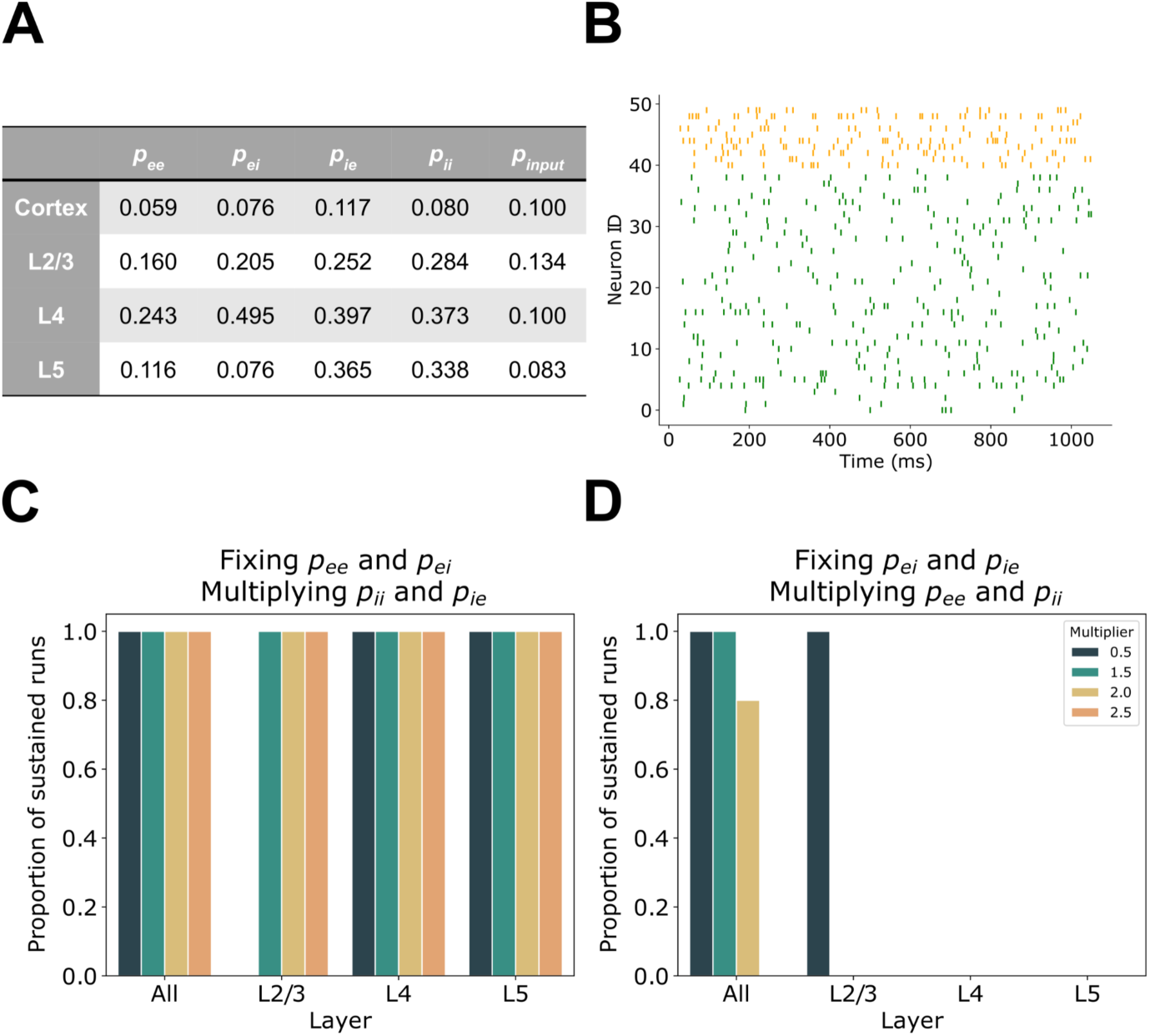
Testing correlated ratios using experimental parameters. (A) Coarse grained connectivity probabilities calculated from Billeh et al. [32]. (B) Raster plot showing the timing of spikes of 50 example neurons in the network with connectivity probabilities experimentally derived from L2/3 using continuous input. Neurons 0-39 (green) are excitatory and neurons 40-49 (orange) are inhibitory. (C) Networks that maintained highly correlated ratios regardless of the value of two parameters showed sustained activity. (D) Same as (C) but instead maintaining weakly correlated ratios which did not result in sustained activity in most trials.

Grid search in the continuous input condition found that the ratios 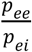 and 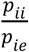 have a strong negative correlation despite not sharing any parameter, while 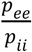 and 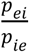, which also have distinct pairs, are not correlated (Fig 2L). We found that the coarse grained experimentally derived connectivity values were consistent with the correlations that we found. We tested the importance of each of these pairs of ratios in sustaining network activity using the connectivity parameters previously reported [32] and found that if the ratios 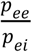 and 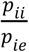 were maintained while changing the actual parameter values, the majority of the resulting networks continued to exhibit sustained activity regardless of how large the values of the probabilities became (up to 0.99; Fig 3C). However, it should not come as a surprise that not all the resulting networks exhibited sustained activity since other potentially critical parameter combinations were being varied simultaneously. In contrast, using those same networks, maintaining the ratios 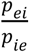 and 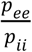 (which are very weakly correlated) did not result in networks that sustained activity even with continuous input (Fig 3D).

### All four classes of connectivity contribute to stiff dimensions

To determine the contribution of individual parameters and their combinations to the stiff dimensions of synaptic architectures, as well as the impact of the input on the model’s stiffness, we used the Fisher Information Matrix (FIM) [15]. By its relation to the Hessian matrix, the FIM evaluated at a specific point in the parameter space examines how the likelihood of matching the spiking activity of murine visual cortex changes along the different dimensions around that point. Large changes in the likelihood along certain dimensions indicates that those dimensions are stiff whereas the others are sloppy. For each class of input, the FIM was computed at the parameter combinations that resulted in the lowest firing rates among the rate-matched networks (8-15 Hz), which we defined as the “optimal” combinations (Fig 4A). For parsimony we selected the lowest rate values since low rates were consistently more difficult to achieve yet revealed generalizable stiff combinations in the large-scale grid search described above. In the grid search, we ran five trials for each parameter combination and found negligible differences in the average firing rates of the networks at the lower end of the target range (8-15 Hz). We therefore estimated the FIM at the five parameter combinations with the lowest average firing rates for each class of inputs.

**Fig 4.**
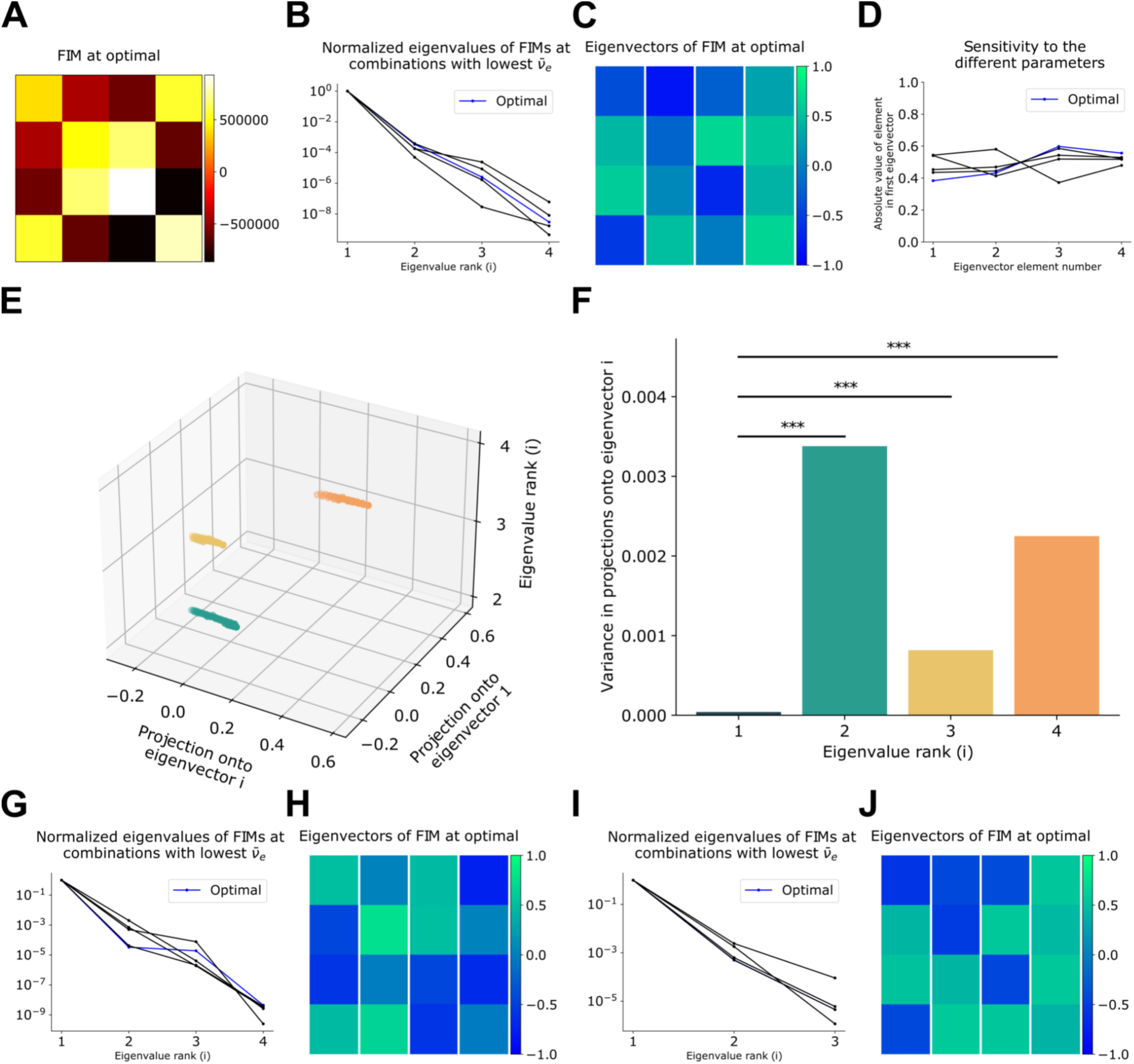
Fisher Information Matrix analysis. (A) FIM computed at the optimal parameter combination for brief input. (B) Eigenvalues of the FIMs computed at the five parameter combinations with the lowest firing rates in the target range using brief input. (C) Eigenvectors of the FIM in A. (D) Sensitivity of the eigenvectors in C to each of the parameters based on the absolute value of each vector element. (E) Projections of parameter vectors that resulted in sustained activity following brief input onto the eigenvectors of the optimal FIM (green: projection onto the eigenvectors 1 and 2; yellow: projection onto the eigenvectors 1 and 3; orange: projection onto the eigenvectors 1 and 4). (F) Variances in the projections in E (*p*_1−2_ ≈ 1.1 ⋅ 10^−14^, *p*_1−3_ ≈ 6.2 ⋅ 10^−10^, *p*_1−4_ ≈ 4.8 ⋅ 10^−14^). (G)&(H) and (I)&(J) As (B)&(C) but for cyclical and continuous input respectively.

To identify the parameter combinations that had the greatest impact on spiking activity, we decomposed the FIM into eigenvectors and identified the corresponding eigenvalues. The eigenvalues of all FIMs extended over several orders of magnitude, consistent with many synaptic architecture parameter combinations being sloppy (Figs 4B, 4G, and 4I). The eigenvectors that corresponded to the largest eigenvalues define the stiffest dimensions and indicated that those specific parameter combinations have the greatest impact on SNN spiking activity. Indeed, when projecting all the parameter vectors with sustained activity onto the different eigenvectors, we find that the first eigenvector— corresponding to the largest eigenvalue—had the lowest variance in projections (Figs 4E and 4F) confirming that the first eigenvector defines the stiffest dimension of parameter space.

We then evaluated the sparsity of the FIM using the Gini coefficient to establish the complexity of the stiff dimensions [20]. In the majority of networks, the Gini coefficient was low (0.21±0.07), indicating that the FIMs were not sparse [20]. The lack of sparsity indicates that the stiff dimensions depend on complex combinations of several parameters and not only on a few critical parameters [15,20], consistent with the results of the grid search. Indeed, the contribution of each parameter to the first eigenvectors reveals that those dimensions are equally sensitive to changes in all of the parameters (Figs 4D, 4H, and 4J). These results were consistent across the three types of inputs.

### Input brevity increasingly restricts the viable wiring parameter space

All the networks that had sustained activity in response to brief input also showed sustained activity when receiving continuous input. In addition, similar patterns of correlations were observed among pairs of parameters for all three types of inputs. For this reason, we hypothesized that dimensions deemed stiff for one case may also be informative of stiff dimensions for other inputs. Indeed, when considering the projections of parameter combinations that resulted in sustained activity using continuous input onto the stiffest dimensions, we found that they are grouped in half the parameter space with little overlap with those networks that didn’t sustain their activity. Importantly, SNNs that demonstrated self-sustaining activity following brief input correspond to a restricted region of the parameter space described by those same eigenvectors (Fig 5A). Interestingly, networks that show sustained activity when receiving cyclical input define an intermediate region of this parameter space as compared with the parameter space defined by networks receiving continuous input, but also encompassing the restricted region of parameter space corresponding with networks receiving brief input (Fig 5A).

**Fig 5.**
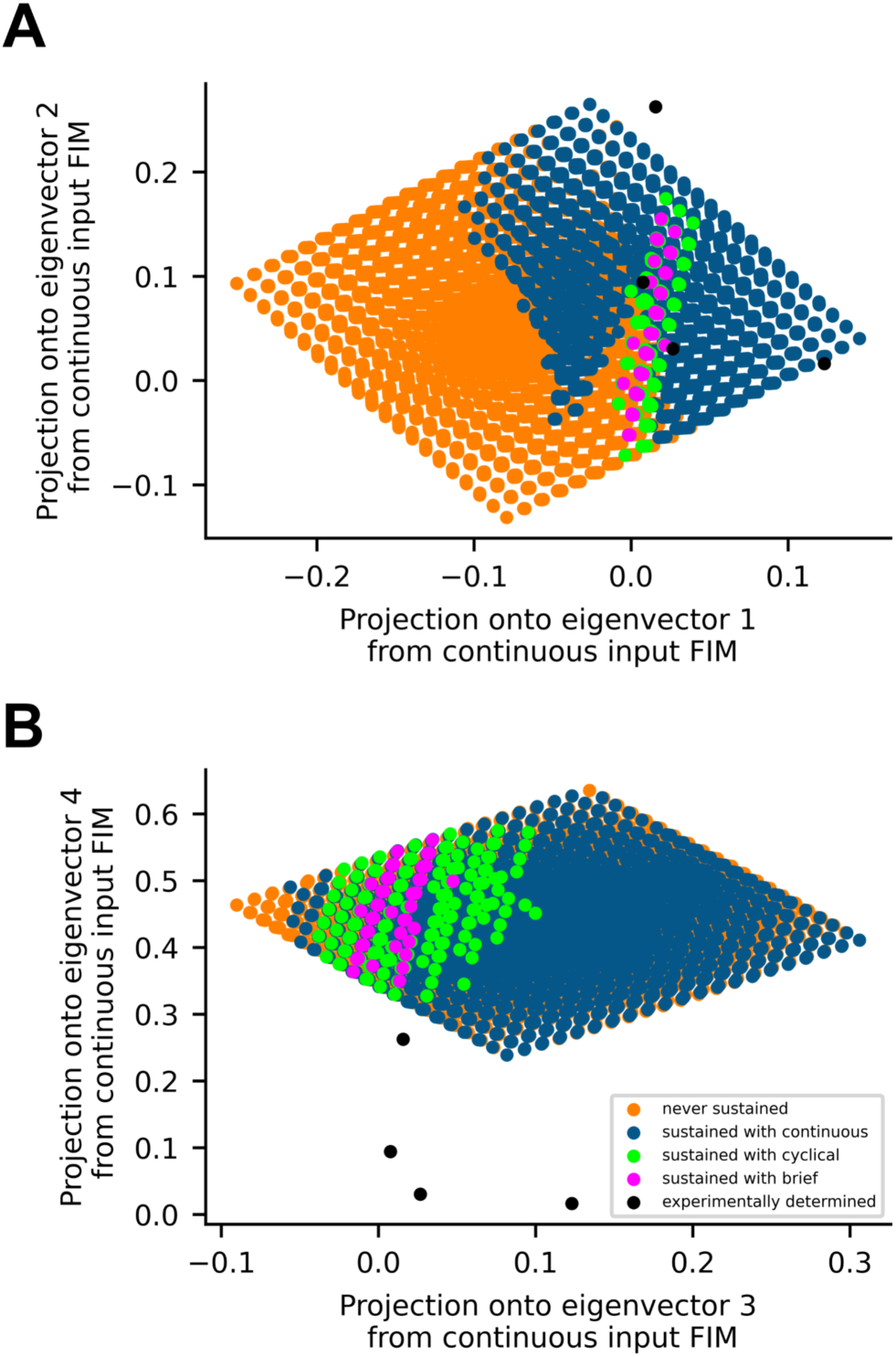
Restrictions of the parameter space along the stiffest dimensions. (A)&(B) Projections of all parameter combinations tested onto the stiffest and sloppiest dimensions respectively, color-coded depending on the resulting spiking activity using each type of input.

Parameter combinations derived from Billeh et al. [32] for L2/3 and for the laminar-agnostic primary visual cortex fall inside the smallest region. However, connectivity parameters from L4 and L5 fall outside of this restricted region and also exhibit sustained activity in response to continuous input (Fig 5A). Notably for L5 parameters, the projection onto the first eigenvector fell in the same very narrow range but the projection onto the second eigenvector differed.

Dimensions used for this analysis define the stiffest dimensions of the system—and consistently, the same type of analysis on the less informative dimensions revealed no structure in the data (Fig 5B).

The non-sparsity of the FIMs indicated the dependence of the stiff dimensions on multiple parameters. We found that the projections of the parameter combinations that exhibited sustained activity following brief input onto the stiffest dimension all fell around 0 (Fig 5A). The eigenvector considered here is e_1_ ≅ [-0.59, 0.43, 0.54, -0.43]. Thus 1.4*p*_*ee*_ − *p*_*ei*_ ≈ −(*p*_*ii*_ − 1.3*p*_*ie*_). This explains the strong negative correlation found between *p*_*ee*_ − *p*_*ei*_ and *p*_*ii*_ − *p*_*ie*_ (Fig 2E). Indeed, the differences and ratios between connectivity probabilities could have been used as the FIM parameters to identify stiff dimensions that depend on only a subset of the parameters considered. However, the results of the grid search indicated that this is not possible because of the strong correlations found between differences and ratios (Fig 2). Hence, inherent relations between the different probabilities of connectivity make it impossible to get stiff dimensions dependent on unique model parameters resulting from straightforward combinations of probabilities of connectivity. Well-founded new methods of describing the connections between neurons at the network level instead of probabilities of connectivity between pairs of neurons could potentially address this problem. For instance, instead of looking at the probability of pairwise connections, we can try considering the probability of different triplet motifs that include both excitatory and inhibitory neurons which might result in less complex stiff dimensions. In the case of neuronal parameters, stiff dimensions that depend on only a subset of the parameters have been obtained [20,21].

## Discussion

Using large scale algorithmic grid searches we found that the parameter space of synaptic architectures is highly anisotropic: large ranges of parameter values produce spiking that matches that of murine visual cortex, while a specific subset of parameter value combinations dramatically change network activity. These are the sloppy and stiff dimensions respectively. Stiff parameter combinations generalize across three broad classes of input into the network. Notably, the region of viable parameter combinations constricted as the requirement for architectures being capable of self-sustaining activity increased.

A recurring theme in neuroscience is that stiff parameter combinations encompass opposing forces. For example, most combinations of ion channel conductances are sloppy with the exception of a maintained ratio between a hyperpolarizing conductance and a depolarizing conductance [17–19,35]. Here we show that the connectivity statistics between and within excitatory and inhibitory neurons comprise the stiff parameter combinations of synaptic architectures. Indeed, maintaining a balance between excitation and inhibition is critical for normal network activity and has been resolved in synaptic conductances at the single cell level *in vivo* [36]. Moreover, many studies have also argued that a balance of excitation and inhibition underlies irregular firing in the neocortex [37,38]. We show that this balance is achieved in synaptic architectures through inter- and intra-population neuronal synaptic connections and involves excitation, inhibition, and disinhibition. It is for this reason that we found that all four connectivity parameters that we implemented contribute to the stiff dimensions and why pairs of connection likelihoods that don’t share any parameters were highly correlated.

Surprisingly, despite simplistic coarse graining, experimentally estimated connectivity probabilities from the entirety of V1 of mice as well as connectivity probabilities from L2/3 of V1 [32] fall inside the small space of topologies that we identified via algorithmic search as potentially capable of self-sustained activity and may suggest a level of autonomy of L2/3 activity not achievable in other laminae. There were notable differences between the algorithmically identified and experimentally measured synaptic architectures in other laminae. Experimentally estimated L5 connectivity parameters themselves did not fall within the restricted space of viable synaptic architectures, although the projection onto the first eigenvector does, indicating congruence with the stiffest dimension. The difference between L5 and algorithmically identified topologies is in the projection onto the second eigenvector, a less stiff dimension, that may reflect differences in the proportions of inhibitory neurons between the laminae. In fact, while the probabilities of connections from subtypes of inhibitory neurons to specific subtypes of neurons do not vary substantially between the different cortical layers [32], the difference we observed in the generalized probabilities is likely due to the predominance of parvalbumin- and somatostatin-positive inhibitory neurons in L5 (48% and 43% of inhibitory neurons respectively) when compared to the percentage of Htr3a-positive neurons (9% vs. 50% in L2/3). As a result, the coarse grained i-i and i-e probabilities in L5 are much higher than the excitatory probabilities in contrast to the other lamina. This could theoretically be compensated for by the much stronger e-e synapses recorded in L5 compared to L2/3 [2,32,39–42]; however, we did not take this particular parameter into consideration in our simulation or analysis. Despite the link between synaptic strength and sustained activity [43], which is directly related to the role of total excitatory drive onto each neuron [44], this is not a parameter that we varied because we opted to focus on the specifics of the synaptic wiring to the extent possible. To do so we used random connectivity weights and conductance-based synapses with time-decaying conductances that result in the total input to each neuron dynamically changing during a single trial since each input is contextualized by the conductance state of the modeled postsynaptic neuron. Notably, it was shown that conductance-based synapses, as compared current-based synapses, result in more informative spiking dynamics [45]. Connectivity parameters of L4 also fall outside the region of the parameter space identified by the stiffest dimensions. This difference lacks an easy explanation but may simply reflect the different roles that layer 4 is hypothesized to play as compared to other lamina and the fact that thalamocortical connectivity is particularly important and heterogeneous into L4 [46].

Machine learning techniques demonstrate that the structure of neural networks can be modified to achieve a specific task and that, following training, these models are capable of accurately modeling neocortical neuronal activity at different stages of the visual processing hierarchy [47]. Here we studied synaptic network architectures identified using first order the spiking statistics rather than trained on a task. It will be of great interest to evaluate whether training similar networks [48] will result in convergence to the same set of synaptic architectures.

Our approach to identify synaptic architectures based solely on spiking statistics and the correspondence with experimental measures demonstrates the utility of spiking statistics as a parsimonious way of studying structure-function relations. Indeed, previous work using maximum entropy models fit to spiking data was similarly successful at identifying cellular-level stiff and sloppy dimensions [20,21]. These results are consistent with a previous experimental study that identified neurons of varying levels of correlation with the network activity [21,49]. Our model on the other hand relied on much simpler and more interpretable descriptives of neuronal firing and elucidated the role of tractable connectivity parameters instead of neuronal ones. In sum, these studies along with this work form a compelling argument to study mesoscale connectivity as well as network wide correlations by fitting models to spiking statistics. In addition, our findings show that coarse-graining of all the different cell types is sufficient for studies of neuronal spiking patterns alone as compared to more complex cortical functions for which these elaborations and more complicated models might become more justified.

## Methods

### Model architecture

Networks consisted of 5000 neurons: 4000 excitatory (*e*) and 1000 inhibitory (*i*). Neurons were modeled as adaptive exponential leaky integrate-and-fire (AdEx) [27] units. Membrane potential of each neuron was governed by the following equations:

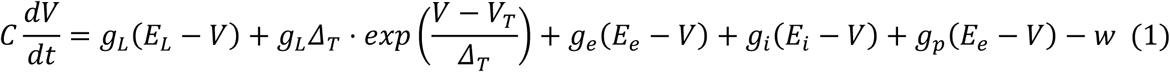

With the decaying adaptation current:

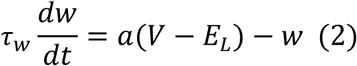

Neurons were connected according to four probabilities of connectivity that were varied: *p*_*ee*_, *p*_*ie*_, *p*_*ei*_,and *p*_*ii*_; where the first subscript index represents the presynaptic neuron type and the second represents the postsynaptic neuron type.

Synaptic conductances decayed exponentially as per the following equations:

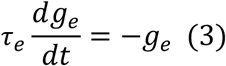

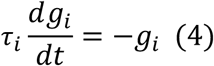

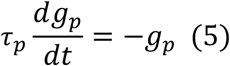

When a neuron fires, the membrane potential is reset, the adaptation current is increased by a value of *b*, and the corresponding synaptic conductance at the synapses receiving the signal is increased by the weight of the connection. The weights of the connections were randomly drawn from a log-normal distribution with the parameters of the corresponding normal being *μ* = −0.64 and *σ* = 0.51. Connections from inhibitory neurons were enhanced by an order of magnitude. The parameters fixed across all simulations are defined and summarized in Table 1.

**Table 1.** Fixed neuronal parameters.

| Parameter Name | Notation | Value |
| --- | --- | --- |
| Capacitance | $C$ | 281 pF |
| Leak conductance | $g_L$ | 30 nS |
| Leak reversal potential | $E_L$ | -70.6 mV |
| Slope factor | $\Delta_T$ | 2 mV |
| Firing threshold | $V_T$ | -40.4 mV |
| Excitatory reversal potential | $E_e$ | 0 mV |
| Inhibitory reversal potential | $E_i$ | -75 mV |
| Excitatory synaptic time constant | $\tau_e$ | 10 ms |
| Inhibitory synaptic time constant | $\tau_i$ | 3 ms |
| Input synaptic time constant | $\tau_p$ | 10 ms |
| Adaptation time constant | $\tau_w$ | 144 ms |
| Subthreshold adaptation | $a$ | 4 nS |
| Spike-triggered adaptation | $b$ | 80.5 pA |

Initial voltages of all neurons were randomly drawn from a normal distribution with mean *μ* = −65*mV* and standard deviation *σ* = 5*mV*.

All simulations were implemented in Python 3 using the Brian Simulator (version 2.2.1) [50].

### Network input

Network activity was initiated using a population of 3000 Poisson units which were connected to both the excitatory and inhibitory neurons in the main network with a connection probability of 0.1 unless otherwise specified, and the weights of the connections were randomly drawn from a log-normal distribution with the parameters of the corresponding normal being *μ* = −0.64 and *σ* = 0.51. Three types of input were considered:

- Brief input: the firing rates of the Poisson units were drawn from a log-normal distribution with the parameters of the corresponding normal being *μ* = 2.8 and *σ* = 0.3, resulting in a mode at around 15 Hz. Their activity was halted after 300 ms.
- Continuous input: similar to the brief input but the activity of the input units was maintained for the duration of the simulation.
- Cyclical input: the maximum firing rates (*v*_*max*_) of the input units were drawn from the same log-normal distribution as that of the brief input, but the instantaneous firing rates varied between 0 and *v*_*max*_ according to a sinusoidal function with a period of 600ms.

### Connectivity parameters

To minimize sampling bias, the four connectivity parameters were algorithmically determined using a low-resolution four-dimensional grid search in which *p*_*ee*_, *p*_*ei*_, and *p*_*ie*_varied between 0.10 and 0.30 and *p*_*ii*_varied between 0.20 and 0.40 with an increment of 0.02. This resulted in 14641 parameter combinations which were each simulated 5 times with different initial membrane potentials, yielding a total of 73205 simulations.

We also tested values based on the experimental data in V1 published by the Allen Institute for Brain Science at an intersomatic distance of 75*μm* [32]. The connectivity statistics reported in the paper are for pyramidal excitatory neurons and the three major classes of inhibitory neurons determined by the markers parvalbumin, somatostatin, and the ionotropic serotonin receptor 5HT3a in all 6 layers of V1. We calculated the four summary connectivity parameters based on the total number of each neuron type also reported in the paper using the following equation:

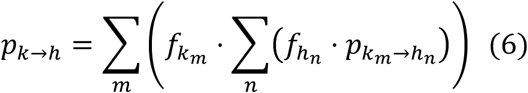

Where:

- *k, h* ∈ {*e, i*};
- *p*_*k*→*h*_ and 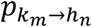 are respectively the connection probability from neurons of type *k* to neurons of type *h*, and the connection probability from neurons of subtype *k*_*m*_to neurons of subtype *h*_*n*_;
- 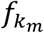 and 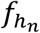 are the fraction of neurons of subtype *k*_*m*_ out of all the *k* neurons, and the fraction of neurons of subtype *h*_*n*_ out of the *h* neurons respectively.

For simulations using the probabilities calculated from the statistics of either L2/3 or L5, the connectivity probability of the input units to the network was computed based on the connections of L4 and L2/3 excitatory neurons respectively to each layer. In the case of L2/3, the resulting value was decreased to account for the numerous inhibitory connections from L4 to excitatory neurons in L2/3.

### Matching spiking statistics to murine visual cortex

The fit of network activity to naturalistic spiking statistics was evaluated based on the following criteria:

- Firing rate of excitatory (*v*_*e*_) and inhibitory (*v*_*i*_) neurons.
- Synchrony (*s*): On runs which were tested individually, synchrony was determined using the Van Rossum distance between excitatory spike trains with 10ms time constant using the “elephant” package [51,52]. For the Fisher Information Matrix calculations (see next subsection), we used the Van Rossum distance on the excitatory spikes during the last 150ms only, since these are representative of the activity of the network after cessation of input and it allows faster computation. However, because computing the Van Rossum distance is computationally expensive, we devised a fast method of estimating the synchrony of the excitatory neurons to use during the grid search: activity was divided into 10ms time bins, and for each time bin we calculated the variance of the number of spikes per neuron divided by the mean number of spikes per neuron and then averaged the result over all time bins. To evaluate the validity of this score, we calculated both this new synchrony measure and the Van Rossum distance for 45 networks that showed sustained activity. The two scores had a 94% correlation.
- Proportion of runs that resulted in sustained activity (*p*_*sus*_): activity is considered sustained if excitatory neurons are firing until the end of the simulation with no more than 150ms of inactivity across the network. In general, networks that showed sustained activity for 1 second sustained their activity for the duration of the simulation regardless of simulation time. However, for runs initiated using brief input, some networks had activity that truncated after 1 second of sustained activity. To ensure that all networks that we evaluated would sustain activity for any simulation duration, we developed an additional check. We selected 50 parameter combinations that resulted in activity sustained for 1 second in at least one trial. We then ran 10 additional trials—each with a new adjacency matrix, initial voltages, and input units—on all 50 parameter combinations. Out of these 500 runs, 435 showed sustained activity for more than 1 second. Of these 435, 85 runs had activity that truncated past one second. We then used this data to train a Support Vector Machine (SVM) classifier with a Radial Basis Function (RBF) kernel to predict which of runs that sustained for 1 second will truncate later. This prediction was based on the excitatory and inhibitory firing rates and the fast synchrony measure calculated on the spikes during the first second of activity only. The best hyperparameters were determined using a cross-validation grid search carried out across different methods of scaling the three features considered. The highest accuracy (96%) was obtained with a power transformation of the scores for a more Gaussian-like distribution and using *Γ* = 0.1 and *C* = 1000. This classifier was then used during the grid search on the connectivity parameters. The SVM was implemented using scikit-learn version 0.22.1 [53]. In this way we ensured that all models considered would spike for the full duration of simulation regardless of the duration.

### Fisher Information Matrix estimation

The likelihood model is a multivariate normal distribution defined by:

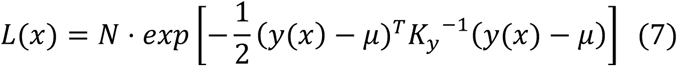

where *N* is a normalization factor; *x* = (*p*_*ee*_, *p*_*ei*_, *p*_*ie*_, *p*_*ii*_) is the vector of model parameters; 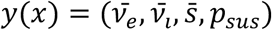 is the vector of simulation scores’ averages from 5 simulations using the same parameter vector *x*; *K*_*y*_ is the (estimated) covariance matrix of the scores vector *y* (estimated using the grid search results); and *μ* is the mean score vector of the distribution. *μ* was set to be equal to the average score vector *y*(*x*) obtained at the optimal parameter combination *x*_*opt*_. As such, *x*_*opt*_ results in the maximum likelihood.

The Fisher Information Matrix (FIM) is equal to the negative of the expected value of the Hessian matrix of the log-likelihood. Consequently, the observed FIM can be estimated as the negative of the Hessian of the log-likelihood [54]. However, computing the Hessian matrix numerically may result in errors, especially if the estimated covariance matrix has bad conditioning. Instead, we derived the expression of the components of the Hessian matrix:

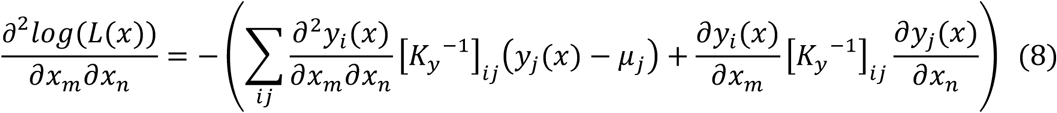

The FIM was computed at the optimal parameter combination which nullifies all the second partial-derivative terms since (*y*(*x*) − *μ*) = 0. We are left with the gradients of the scores which were estimated by calculating all four scores for networks that have 3 of the connectivity parameters fixed and one of them varied between *x*_*optimal*_ − 0.01 and *x*_*optimal*_ + 0.01, then fitting a line to these results due to the strong linear relationships around the optimal values (Fig 6).

**Fig 6.**
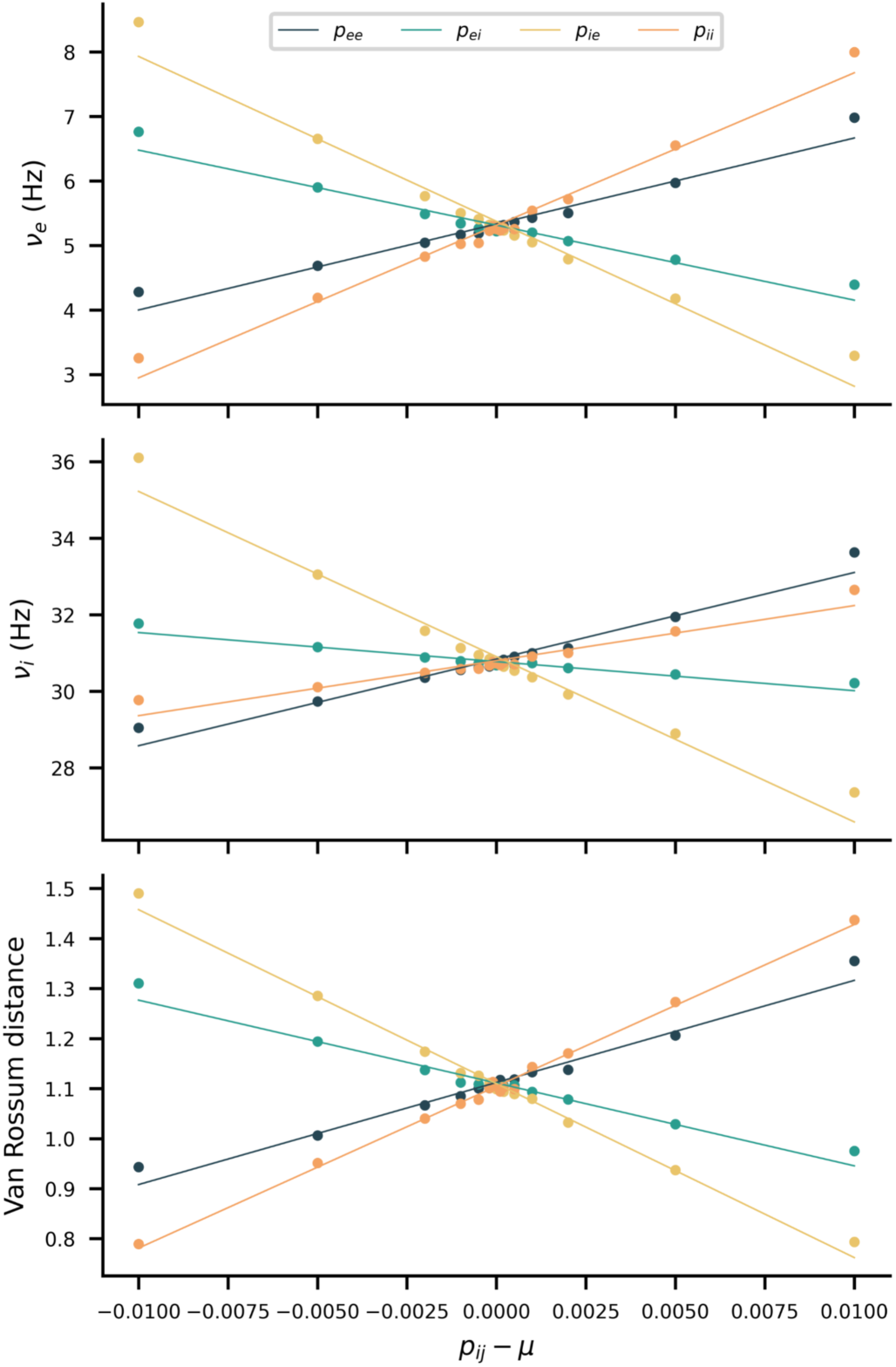
Variation of each spiking activity score with the change in each probability of connectivity around the optimal points.

### Gini Coefficient

Sparsity of matrices was evaluated with the Gini coefficient [55] using the code provided by Panas et al. [20]:

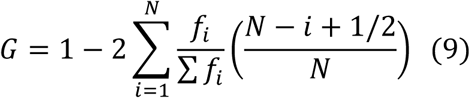

Where *f*_*i*_ are the values of the *N* elements in the matrix.

### Statistical testing

Variances in the projections of the parameter vectors onto the eigenvectors were compared using the Levene test.

## Acknowledgements

We thank former and current MacLean lab members, Yuqing Zhu, Isabel Garon, Maayan Levy, and Gabriella Wheeler Fox, and Dr. Emil Sidky for assistance with initial simulations, accurate estimations of Fisher Information Matrices, and helpful comments on our manuscript.

